# Helmsman: fast and efficient generation of input matrices for mutation signature analysis

**DOI:** 10.1101/373076

**Authors:** Jedidiah Carlson, Jun Z Li, Sebastian Zöllner

## Abstract

**Motivation:** The spectrum of somatic single-nucleotide variants in cancer genomes often reflects the signatures of multiple distinct mutational processes, which can provide clinically actionable insights into cancer etiology. Existing software tools for identifying and evaluating these mutational signatures do not scale to analyze large datasets containing thousands of individuals or millions of variants.

**Results:** We introduce *Helmsman*, a program designed to rapidly generate mutation spectra matrices from arbitrarily large datasets. *Helmsman* is up to 300 times faster than existing methods and can provide more than a 100-fold reduction in memory usage, making mutation signature analysis tractable for any collection of single nucleotide variants, no matter how large.

**Availability:** *Helmsman* is freely available for download at https://github.com/carjed/helmsman under the MIT license. Detailed documentation can be found at https://www.jedidiahcarlson.com/docs/helmsman/, and an interactive Jupyter notebook containing a guided tutorial can be accessed at https://mybinder.org/v2/gh/carjed/helmsman/master.

**Supplementary information:** Supplementary information for this article is available.

## 1 Introduction

The spectrum of somatic single-nucleotide variants (SNVs) in cancer genomes carries important information about the underlying mutation mechanisms, providing insight into the development, evolution, and etiology of the cancer cell populations (Alexandrov, Nik-Zainal, Wedge, Aparicio, *et al.,* 2013). Evaluating these patterns of variation, broadly referred to as “mutational signatures,” has become an important task in precision oncology, as mutational signatures can be used both to refine cancer diagnoses and identify effective targeted therapies (Kumar-Sinha and Chinnaiyan, 2018).

Several software programs have been developed to aid researchers in identifying and evaluating the mutational signatures present in cancer genomes (Gehring *et al.,* 2015; Rosenthal *et al.,* 2016; Rosales *et al.,* 2017). Most methods consider 96 mutation subtypes, defined by the type of base change (C>A, C>G, C>T, T>A, T>C, T>G) and the trinucleotide sequence context (e.g., C[<u>T</u>><u>G]</u>T, C[<u>C</u>><u>A]</u>T, and so on) (Alexandrov, Nik-Zainal, Wedge, Campbell, *et al.,* 2013). Mutation signature analysis methods aim to express the observed mutation spectrum in each sample as a linear combination of K distinct mutational signatures, where the signatures are inferred directly from the input data, or taken from external sources such as the COSMIC mutational signature database (Alexandrov, Nik-Zainal, Wedge, Aparicio, *et al.,* 2013). These programs typically start with an input file, often in a standard format such as Variant Call Format (VCF) or Mutation Annotation Format (MAF), containing the genomic coordinates of each SNV and the sample(s) in which they occur. As a first step, these SNVs must be summarized into a NxS mutation spectra matrix, M, containing the frequencies of S different SNV subtypes in each of N unique samples (where the *M_i,j_* entry indicates the number of observed SNVs of subtype j in sample i). Most methods are implemented as R packages and must read the entire input file into memory prior to generating the mutation spectra matrix. For large input files, containing for example millions of SNVs and hundreds or thousands of samples, the memory required for this step can easily exceed the physical memory capacity of most servers, rendering such tools incapable of directly analyzing large datasets. To circumvent these computational bottlenecks, researchers must either limit their analyses to small samples, pool samples together, or develop new software to generate the mutation spectra matrix.

## 2 Description

To overcome the limitations of existing mutation signature analysis tools, we have developed a Python application, named *Helmsman*, for rapidly generating mutation spectra matrices from arbitrarily large datasets. *Helmsman* accepts both VCF and MAF files as input.

For each SNV in a VCF file, *Helmsman* identifies the mutation type based on the reference and alternative alleles, then queries the corresponding reference genome for the trinucleotide context of the SNV, determining subtype j. The genotypes of the N samples for this SNV are represented as an integer array, with the number of alternative alleles per sample coded as 0, 1, or 2 according to the observed genotype (Pedersen and Quinlan, 2017). *Helmsman* then updates the jth column of the mutation spectra matrix by vectorized addition of the genotype array (i.e., *M_ij_* is incremented by 1 if individual *i* is heterozygous but does not change if individual *i* is homozygous for the reference allele). Consequently, *Helmsman’s* processing time is independent of sample size and scales linearly with the number of SNVs. The only objects stored in memory are the array of N genotypes for the SNV being processed and the Nx96 mutation spectra matrix, so memory usage is independent of the number of SNVs and scales linearly with sample size.

## 2.1 Additional Features

In addition to being optimized for speed and low memory usage, *Helmsman* includes several features to accommodate various usage scenarios and minimize the amount of pre-processing necessary to analyze large mutation datasets. For example, if input data are spread across multiple files (e.g., by different sub-samples or genomic region), *Helmsman* can process these files in parallel and aggregate them into a single mutation spectra matrix, providing additional performance improvements and avoiding the need to generate intermediate files. Similarly, in certain applications, it may be desirable to pool similar samples together (e.g., by tumor type) when generating the mutation spectra matrix. *Helmsman* can pool samples on-the-fly, without needing to pre-annotate or reshape the input file with the desired grouping variable.

*Helmsman* also includes basic functionality for extracting mutation signatures from the mutation spectra matrix using non-negative matrix factorization (NMF) or principal component analysis (PCA) functions from the *nimfa* (Žitnik and Zupan, 2012) and *scikit-learn* (Pedregosa *et al.,* 2011) Python libraries, respectively. Alternatively, *Helmsman* can generate an R script with all code necessary to load the output matrix into R and apply existing supervised and unsupervised mutation signature analysis programs without requiring users to perform the computationally expensive task of generating this matrix from within the R environment. All features are described in detail in the online documentation.

## 3 Results

We compared *Helmsman’s* performance to that of three published R packages: *SomaticSignatures* (Gehring *et al.*, 2015), *deconstructSigs* (Rosenthal *et al.,* 2016), and *signeR* (Rosales *et al.,* 2017). We also considered several other tools and discuss their performance in the **Supplementary Material**. For our tests, we generated a small VCF file (2.7MB compressed with bgzip) containing 15,971 germline SNVs on chromosome 22 from 2,504 samples sequenced in the 1000 Genomes Project phase 3 (1000 Genomes Project Consortium *et al.,* 2015), and measured the runtime and memory usage necessary for each program to generate the mutation spectra matrix. We also attempted to run each program using the full chromosome 22 VCF file from the 1000 Genomes Project, containing 1,055,454 SNVs in 2504 individuals.

All programs generated the same mutation spectra matrices. *Helmsman* processed the small VCF file in 8 seconds, with a memory footprint of 140MB, and the full VCF file in 482 seconds (linear increase for ∼60x more variants) with no increase in memory usage as the sample size remained the same. In contrast, to process the small VCF file, *SomaticSignatures* took 227 seconds with a memory footprint of 18GB, *deconstructSigs* took 2,376 seconds and 7.5GB of memory, and *signeR* took 1,740 seconds and 10.2GB of memory (**Fig. 1**). None of these were able to load the full VCF file due to memory allocation errors. All other tools we considered showed similar performance bottlenecks (**Supplementary Material**; **Supplementary Fig. S1**).

**Fig. 1.**
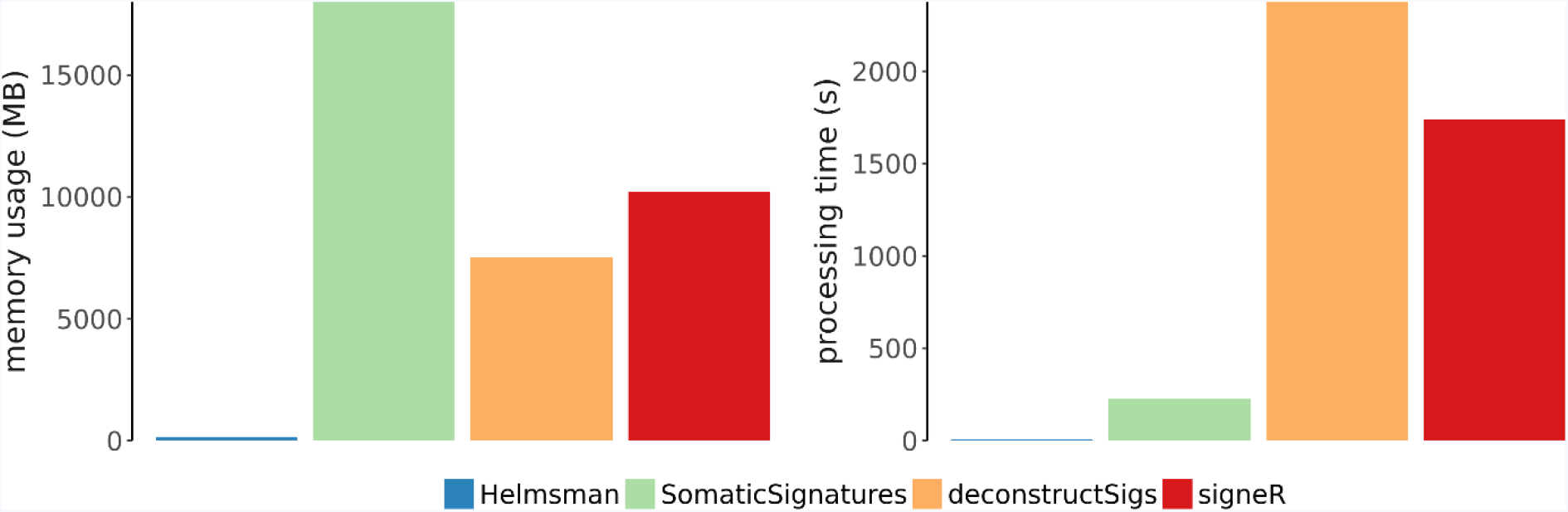
Performance comparison for generation of the mutation spectra matrix from a VCF file containing 15,971 SNVs in 2,504 samples.

To further highlight the speed and efficiency of *Helmsman* for large datasets, we evaluated the entire set of 36,820,990 autosomal biallelic SNVs from the 1000 Genomes phase 3 dataset (14.4 GB when compressed with bgzip). Using 22 CPUs (one per chromosome VCF file), *Helmsman* generated the mutation spectra matrix in 64 minutes (approximately 1.5 seconds per sample), with each process requiring <200MB of memory.

## 4 Discussion

As massive sequencing datasets become increasingly common in areas of cancer genomics and precision oncology, there is a growing need for software tools that scale accordingly and can be integrated into automated workflows. Our program, *Helmsman*, provides an efficient, standardized framework for generating mutation spectra matrices from arbitrarily large, multi-sample VCF or MAF files. For small datasets, *Helmsman* performs this task up to 300 times faster than existing methods and is the only tool that can be directly applied to modern large sequencing datasets.

